# Generation of a new *six1*-null line in *Xenopus tropicalis* for study of development and congenital disease

**DOI:** 10.1101/2021.07.30.454496

**Authors:** Kelsey Coppenrath, Andre L.P. Tavares, Nikko-Ideen Shaidani, Marcin Wlizla, Sally A. Moody, Marko Horb

## Abstract

The vertebrate Six (<u>S</u>ine oculis homeobo<u>x</u>) family of homeodomain transcription factors play critical roles in the development of several organs. Six1 plays a central role in cranial placode development, including the precursor tissues of the inner ear, as well as other cranial sensory organs and the kidney. In humans, mutations in *SIX1* underlie some cases of branchio-oto-renal syndrome (BOR), which is characterized by moderate to severe hearing loss. We utilized CRISPR/Cas9 technology to establish a *six1* mutant line in *Xenopus tropicalis* that is available to the research community. We demonstrate that at larval stages, the *six1-null* animals show severe disruptions in gene expression of putative Six1 target genes in the otic vesicle, cranial ganglia, branchial arch and neural tube. At tadpole stages, *six1-null* animals display dysmorphic Meckel’s, ceratohyal and otic capsule cartilage morphology. This mutant line will be of value for the study of the development of several organs as well as congenital syndromes that involve these tissues.

## COMBINED RESULTS AND DISCUSSION

The vertebrate Six (<u>S</u>ine oculis homeobo<u>x</u>) family of homeodomain transcription factors is comprised of 6 genes (*Six1*-*Six6*) that are highly related to 3 *Drosophila* genes: *sine oculis (so), optix*, and *six4* (Kawakami et al., 2000; Kumar, 2009). Six1 and Six2 are highly related to *so*, which is a key regulator of the fly visual system, but in vertebrates *Six1* and *Six2* play critical roles in the development of several organs including muscle, thymus, kidney, lung and mammary gland (Laclef et al., 2003; Grifone et al., 2005; Kobayashi et al., 2007; Coletta et al., 2010; El-Hashash et al., 2011). Several studies demonstrate that Six1 has a central role in cranial placode development. Six1 loss-of-function in *Xenopus* and chick results in reduced expression of several placode genes and defects in otic development (Brugmann et al., 2004; Schlosser et al., 2008; Christophorou et al., 2009). In zebrafish, Six1 knock-down results in loss of inner ear hair cells (Bricaud and Collazo, 2006). *Six1-*null mice show defects in the olfactory placode, inner ear and cranial sensory ganglia (Ikeda et al., 2007; Konishi et al., 2006; Laclef et al., 2003; Ozaki et al., 2004; Zou et al., 2004). Thus, Six1 is a critical gene for the normal development of a number of organs.

Six1 also has been identified as one of the underlying genetic causes of Branchio-oto-renal syndrome (BOR) (OMIM 608389). BOR is the second most common type of autosomal dominant syndrome with hearing loss (Smith, 2018), and is included in genetic screens for hearing loss, such as the OtoSCOPE panel at the University of Iowa (https://morl.lab.uiowa.edu/otoscope%C2%AE-genetic-hearing-loss-testing-v9) and the Branchio-oto-renal Syndrome panel at Cincinnati Children’s Hospital (https://www.cincinnatichildrens.org/service/g/genetic-hearing-loss/requisition). Mutations in SIX1 occur in the N-terminal protein-protein interaction domain (SD) or in the DNA-binding homeodomain (HD). A recent study also found Six1 mutations in patients with craniosynostosis and hearing defects (Calpena et al., 2021). Because Six1 has important roles in both normal and abnormal development of many tissues, we used Crispr/Cas 9 technology to create a null line in *Xenopus tropicalis*, an animal model highly suited for studying the cell and molecular regulation of gene function (Tandon et al 2017, Kakebeen and Wills 2019, Bhattacharya et al 2015).

### Generation of the line

To generate a *six1* knockout, we designed two sgRNAs to the first exon of *six1* gene; mutations in this region would disrupt the SD domain. Injection of either sgRNA with Cas9 resulted in small, stunted tadpoles and eventual embryonic lethality before stage 47 (data not shown). To overcome embryonic lethality, we injected the first sgRNA into all eight vegetal blastomeres at the 32-cell stage. This approach would localize the sgRNAs and Cas9 to prospective endodermal cells where the germ cells are located and avoid creating mutations in neural and mesoderm, where *six1* is expressed. Seven F0 founders survived to adulthood and once they reached sexual maturity, they were outcrossed to wild type to generate F1 offspring. Of these seven F0 founders, three (two female and one male) produced offspring with germline mutations; two males did not produce offspring, two females showed no germline transmission. Initial genotyping of F1 embryos contained +1bp and -4bp mutations, but none survived; female 2 offspring had +8bp and +1bp mutations; male 1 offspring had a +1bp mutation. We raised sibling embryos from female 2 and male 1 to adulthood and genotyped them as adults. Of 26 F1 adults from female 1, we identified one with a -28bp mutation, fourteen with a +1bp mutation, three with a -9bp mutation, one with a -6bp mutation and one with a +8bp mutation. We then generated more -28bp heterozygous F2 *six1* mutants by outcrossing the F1 male to wild type. F2 -28/+ *six1* mutants were then intercrossed to generate F3 mutants that were used for phenotypic characterization in this study. This -28bp deletion leads to a frameshift at amino acid 53 resulting in a stop codon 36 amino acids downstream (Figure 1).

**Figure 1:**
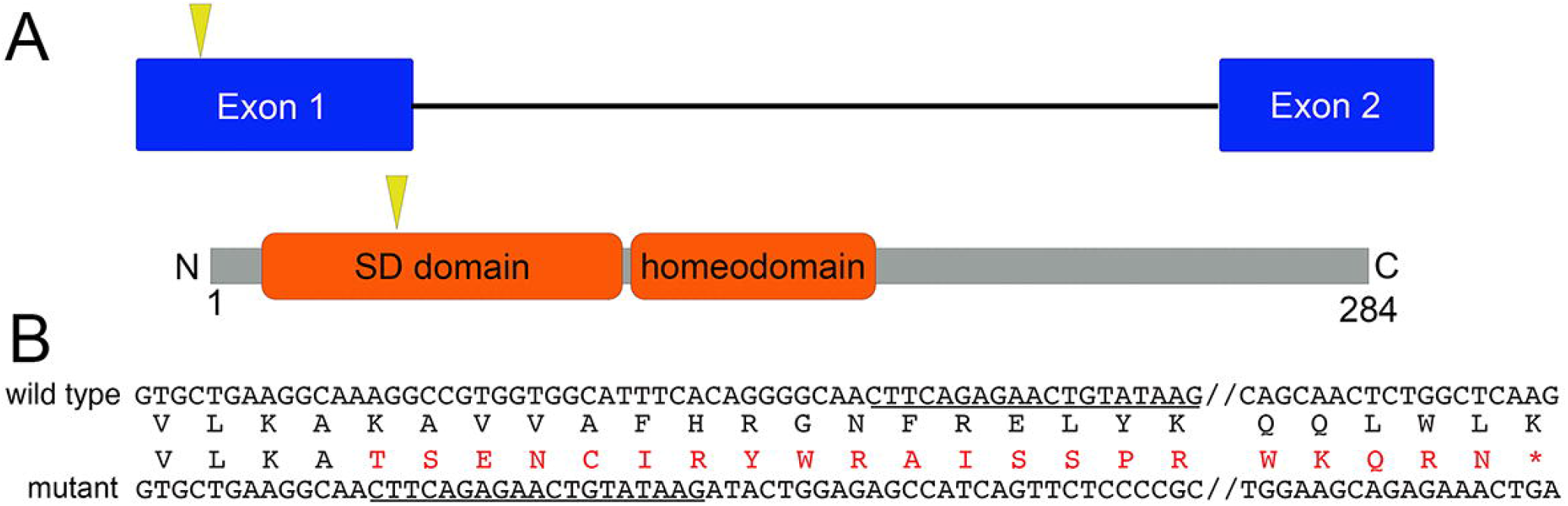
Generation of *six1* mutant *X. tropicalis* line. A. Schematic outline of genomic *six1* locus and Six1 protein. Yellow arrowhead demarcates the relative location of sgRNA target sequence and mutations in this region are expected to disrupt the protein-protein interaction domain (SD). B. Alignment of wild type and *Xtr*.*six1*^*em2Horb*^ mutant nucleotide sequence with the -28bp deletion. Underlined region defines where the mutant and wild type sequence are identical after the 28bp deletion. Amino acid sequence is included to show frameshift with resulting stop codon 36 amino acids downstream.

### Six1 otic targets are differentially affected by loss of Six1

A number of large-scale screens in flies and vertebrates have identified hundreds of potential SO/Six1 transcriptional targets in numerous tissues at several developmental stages (Ando et al., 2005; Jusiak et al., 2014; Yan et al., 2015; Riddiford and Schlosser, 2016). To assess the effects of loss of Six1 in the otic vesicle (OV), which is known to require Six1 for proper development (Ozaki et al., 2004; Zou et al., 2004), we chose to investigate the expression of genes that, based on a ChIP analysis of the E10.5 mouse OV (Li et al., 2020), are in close proximity to sites bound by Six1. Using an in situ hybridization assay, we found that some of these putative target genes showed subtle changes in their expression patterns whereas others showed significant loss of expression.

There are 3 Six1-occupied binding sites in proximity to *pax2* in the mouse OV (Li et al., 2020). At *X. laevis and X. tropicalis* larval stages, *pax2* is expressed in the OV, neural tube and ventral retina (Heller and Brandli, 1997; image XB-IMG-2480 in Xenbase (http://www.xenbase.org/, RRID:SCR_003280). There is a similar pattern in *X. tropicalis six1*^*+/-*^ and *six1*^*-/-*^ larvae (Figure 2A; Table 1), indicating Six1 is not required for *pax2* otic expression, consistent with a previous report in mouse (Ozaki et al., 2004). There are 13 Six1-occupied binding sites in proximity to *eya2* in the mouse OV (Li et al., 2020). In larval *X. laevis, eya2* is expressed in the OV, olfactory placode, cranial ganglia, nephric mesoderm and hypaxial muscle precursors (Neilson et al., 2010). There is a similar pattern in wild-type (not shown) and heterozygous *X. tropicalis* (Figure 2B; Table 1). In *six1*-null larvae, the *eya2* expression pattern was differentially perturbed in different tissues: it was not detected in the olfactory placode, IX/X ganglion or hypaxial muscle precursors in some larvae; the OV expression domain was irregular or smaller, and expression in the VII ganglion was thinner, truncated or discontinuous (Figures 2B, 3A-B; Table 1). There is 1 Six1-occupied binding site in proximity to *tbx1* in the mouse OV (Li et al., 2020). In larval *X. tropicalis, tbx1* is expressed in the ventral OV and the branchial arches (BAs) (Showell et al., 2006). There is a similar pattern in every *six1*^*+/-*^ larva (Figures 2C, 3C-D); in some *six1*^*-/-*^ larvae OV and BA staining was less distinct and in others not detected (Figures 2C, 3C-D; Table 1). There is 1 Six1-occupied binding site in proximity to *dlx5* in the mouse OV (Li et al., 2020). In *X. tropicalis six1*^*+/+*^ and *six1*^*+/-*^ larvae, *dlx5* is expressed in the OV, BA and olfactory placode (image XB-IMG-3021 in Xenbase (http://www.xenbase.org/, RRID:SCR_003280; Figure 2D; Table 1). In *six1*^*-/-*^ larvae, *dlx5* staining was not detected in any of these tissues (Figure 2D; Table 1). There are 6 Six1-occupied binding sites in proximity to *irx1* in the mouse OV (Li et al., 2020). In larval *X. tropicalis six1*^*+/+*^ and *six1*^*+/-*^, *irx1* is expressed in the OV and neural tube (Figure 2E; Table 1). A previous study that analyzed the effects of reducing Six1 levels in *Xenopus* embryos by antisense morpholino oligonucleotides or Crispr/Cas9 demonstrated that *irx1* OV expression requires Six1 (Sullivan et al., 2019); this was confirmed in *six1*^*-/-*^ larvae in which OV staining was not detected and neural tube expression was reduced (Figures 2E, 3E; Table 1). There are 4 Six1-occupied binding sites in proximity to *sobp* in the mouse OV (Li et al., 2020). In *X. laevis* and in *X. tropicalis six1*^*+/+*^ and *six1*^*+/-*^ larvae, *sobp* is expressed in the OV, olfactory placode, neural tube, and cranial ganglia (Neilson et al., 2010; Figure 2F); in *six1*^*-/-*^ larvae, *sobp* staining was not detected in any of these structures (Figures 2F, 3F; Table 1). There are 3 Six1-occupied binding sites in proximity to *rnf150* in the mouse OV (Li et al., 2020). In *X. tropicalis six1*^*+/+*^ (not shown) and *six1*^*+/-*^ larvae, *rnf150* is expressed in the OV, BA, neural tube and retina (Figure 2G; Table 1); in *six1*^*-/-*^ larvae, *rnf150* staining was not detected in the OV and was reduced in the other structures (Figures 2G, 3G; Table 1). There is 1 Six1-occupied binding site in proximity to *pick1* in the mouse OV (Li et al., 2020). In *X. tropicalis six1*^*+/+*^ (not shown) and *six1*^*+/-*^ larvae, *pick1* is expressed in the OV, BA, retina and neural tube (Figure 2H; Table 1); in *six1*^*-/-*^ larvae, *pick1* staining was not detected in any of these tissues (Figures 2H, 3H; Table 1).

**Table 1:** Frequency of expression of putative Six1 target genes in various tissues.

|  |  |  |  |  |  |  |
| --- | --- | --- | --- | --- | --- | --- |
| ***pax2*** | **OV** | **NT** | **Retina** |  |  |  |
| *six1*<sup>+/-</sup> | 100% (5) | 100% (5) | 100% (5) |  |  |  |
| *six*<sup>-/-</sup> | 100% (5) | 100% (5) | 100% (5) |  |  |  |
| ***eya2*** | **OV** | **Olf** | **VIIg** | **IX/Xg** | **Hyp** | **Nephric** |
| *six1*<sup>+/-</sup> | 100% (10) | 100% (10) | 100% (10) | 40% (10) | 100% (10) | 100% (10) |
| *six*<sup>-/-</sup> | 100% (4)<br>but smaller,<br>malformed | 50% (4) but<br>reduced | 100% (4)<br>but<br>truncated | 0% (4) | 75% (4) but<br>reduced | 75% (4) but<br>reduced |
| ***tbx1*** | **OV** | **BA** |  |  |  |  |
| *six1*<sup>+/-</sup> | 100% (6) | 100% (6) |  |  |  |  |
| *six*<sup>-/-</sup> | 82% (11) | 64% (11);<br>not distinct |  |  |  |  |
| ***dlx5*** | **OV** | **BA** | **Olf** |  |  |  |
| *six1*<sup>+/-</sup> | 100% (15) | 100% (15) | 100% (15) |  |  |  |
| *six*<sup>-/-</sup> | 0% (8) | 0% (8) | 0% (8) |  |  |  |
| ***irx1*** | **OV** | **NT** |  |  |  |  |
| *six1*<sup>+/-</sup> | 100% (5) | 100% (5) |  |  |  |  |
| *six*<sup>-/-</sup> | 0% (5) | 80% (5) |  |  |  |  |
| ***sobp*** | **OV** | **Olf** | **NT** | **VIIg** |  |  |
| *six1*<sup>+/-</sup> | 100% (15) | 100% (15) | 100% (15) | 100% (15) |  |  |
| *six*<sup>-/-</sup> | 0% (5) | 0% (5) | 100% (5)<br>but reduced | 0% (5) |  |  |
| ***rnf150*** | **OV** | **BA** | **NT** | **Retina** |  |  |
| *six1*<sup>+/-</sup> | 100% (15) | 100% (15) | 100% (15) | 100% (15) |  |  |
| *six*<sup>-/-</sup> | 0% (4) | 100% (4)<br>but reduced | 0% (4) | 0% (4) |  |  |
| ***pick1*** | **OV** | **BA** | **NT** | **Retina** |  |  |
| *six1*<sup>+/-</sup> | 100% (10) | 100% (10) | 100% (10) | 100% (10) |  |  |
| *six*<sup>-/-</sup> | 0% (4) | 0% (4) | 0% (4) | 0% (4) |  |  |

**Legend:** BA, branchial arches; Hyp, hypaxial muscle precursors; Nephric, nephric mesoderm; NT, neural tube; Olf, olfactory placode; OV, otic vesicle; VIIg, facial ganglion placode; IX/Xg, glossopharyngeal/vagal ganglion placode. Numbers in brackets denote number of larvae analyzed.

**Figure 2:**
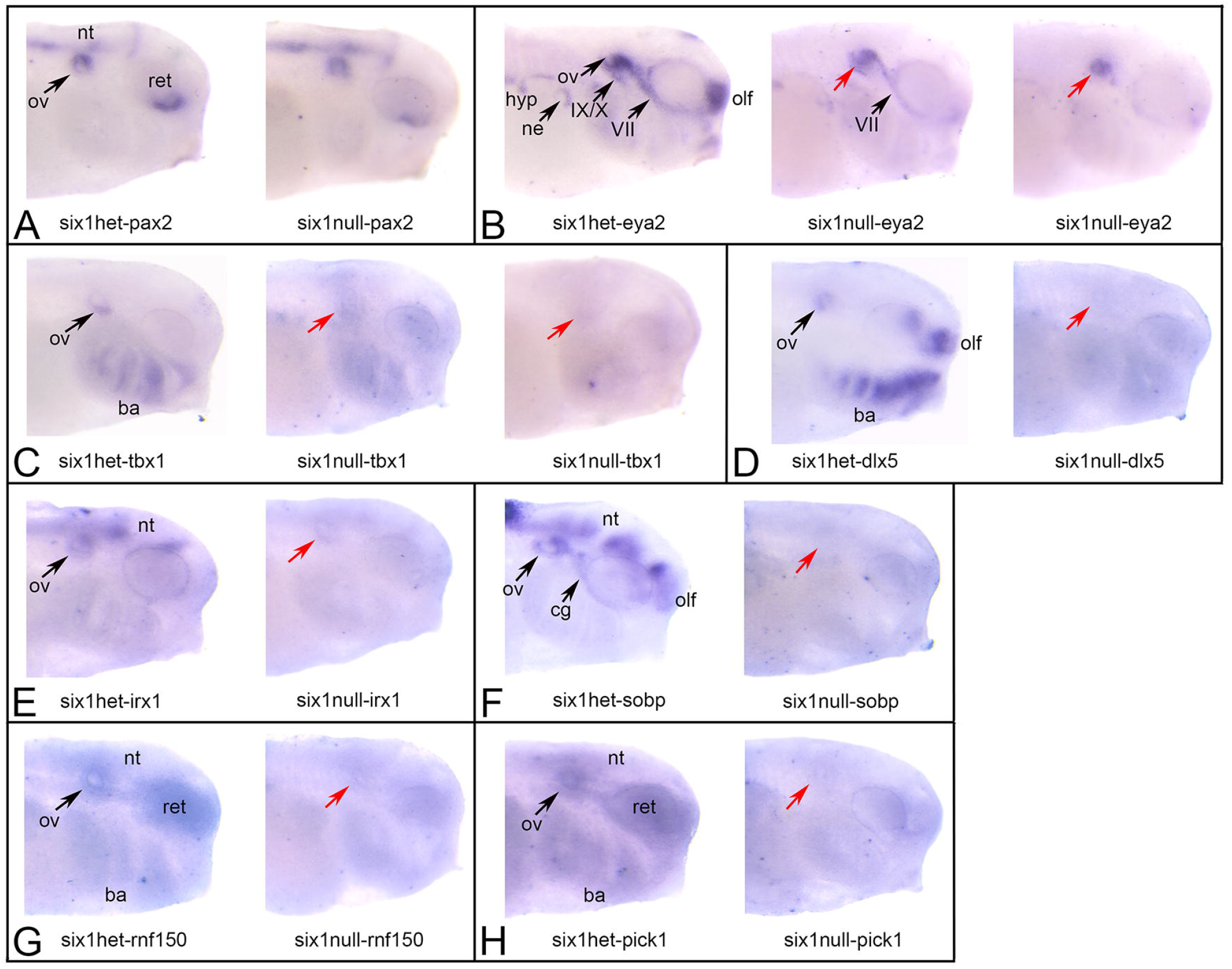
In situ hybridization of *six1*^*+/-*^ and *six1* ^*-/-*^ *X. tropicalis* larvae A. Left image: *pax2* expression in a *six1*^*+/-*^ larva. Staining is strong in the otic vesicle (ov), retina (ret) and neural tube (nt). Right image: *pax2* expression in a *six1*^*-/-*^ larva is indistinguishable from the heterozygous sibling. B. Left image: *eya2* expression in a *six1*^*+/-*^ larva. Staining is strong in ov, olfactory placode (olf), VII and IX/X cranial ganglia, nephric mesoderm (ne) and hypaxial muscle precursors (hyp). Middle image: *eya2* expression in a *six1*^*-/-*^ larva with a moderate effect. ov (red arrow) and VIIg staining appears normal, ne and hyp staining appears reduced, and IX/Xg and olf staining is not detectable. Right image: *eya2* expression in a *six1*^*-/-*^ larva with a severe effect. Only ov staining (red arrow) is detected and ov is smaller than normal. C. Left image: *tbx1* expression in a *six1*^*+/-*^ larva. Staining is strong in ventral ov and branchial arches (ba). Middle image: *tbx1* expression in a *six1*^*-/-*^ larva with a moderate effect. ov (red arrow) and ba staining is reduced. Right image: *tbx1* expression in a *six1*^*-/-*^ larva with a severe effect. ov (red arrow) and ba staining is not detected. D. Left image: *dlx5* expression in a *six1*^*+/-*^ larva. Staining is strong in ov, ba and olf. Right image: Staining in ov (red arrow), ba and olf is not detected. E. Left image: *irx1* expression in a *six1*^*+/-*^ larva. Staining is strong in ov and nt. Right image: *irx1* expression in a *six1*^*-/-*^ larva. Staining in ov (red arrow) and nt is not detected. F. Left image: *sobp* expression in a *six1*^*+/-*^ larva. Staining is strong in ov, VIIg (cg), olf and nt. Right image: *sobp* expression in a *six1*^*-/-*^ larva. Staining in ov (red arrow), cg, olf and nt is not detected. G. Left image: *rnf150* expression in a *six1*^*+/-*^ larva. Staining is strong in ov, ba, ret and nt. Right image: *rnf150* expression in a *six1*^*-/-*^ larva. Staining in ov (red arrow), ret and nt is not detected, and greatly reduced in ba. H. Left image: *pick1* expression in a *six1*^*+/-*^ larva. Staining is strong in ov, ba, ret and nt. Right image: *pick1* expression in a *six1*^*-/-*^ larva. Staining in ov (red arrow), ba, ret and nt is not detected. All views are lateral with anterior to the right and dorsal to the top.

**Figure 3:**
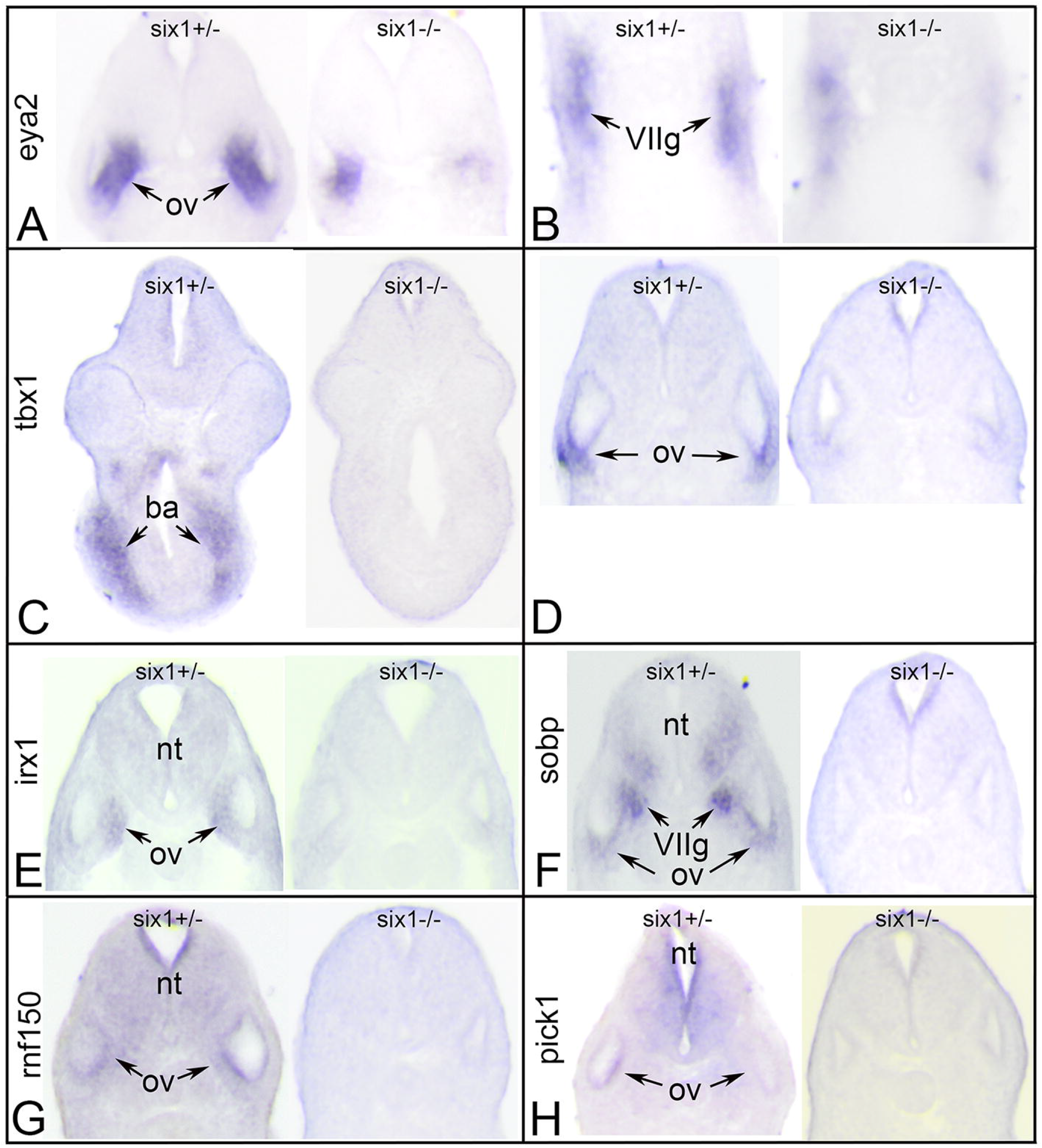
Changes in gene expression after loss of one (*six1*^*+/-*^) or two (*six1*^*-/-*^) alleles of *six1*. A. Sections through the otic vesicle (ov) of *six1* heterozygous (left) and *six1* null larva processed for *eya2* expression. Note the smaller domain of *eya2* expression in the null ov. B. Sections through the facial ganglion placode (VIIg) of *six1* heterozygous (left) and *six1* null larva processed for *eya2* expression. Note the discontinuous nature of *eya2* expression in the null VIIg. C. Sections through the branchial arches (ba) of *six1* heterozygous (left) and *six1* null larva processed for *tbx1* expression. Note the lack of *tbx1* expression in the null ba. D. Sections through the ov of *six1* heterozygous (left) and *six1* null larva processed for *tbx1* expression. Note the lack of *tbx1* expression in the null ov. E. Sections through the ov of *six1* heterozygous (left) and *six1* null larva processed for *irx1* expression. Note the lack of *irx1* expression in the null neural tube (nt) and ov. F. Sections through the ov of *six1* heterozygous (left) and *six1* null larva processed for *sobp* expression. Note the lack of *sobp* expression in the null ventral nt, VIIg and ov. G. Sections through the ov of *six1* heterozygous (left) and *six1* null larva processed for *rnf150* expression. Note the lack of *rnf150* expression in the null nt and ov. H. Sections through the ov of *six1* heterozygous (left) and *six1* null larva processed for *pick1* expression. Note the lack of *pick1* expression in the null nt and ov. All sections are in the transverse plane oriented with dorsal to the top.

Together, these data point out several important features of Six1 regulation of craniofacial genes. First, if a gene expressed in the OV has a Six1-occupied site in close proximity, it is highly likely that Six1 will be required for its expression there. Of the 8 genes assayed, only *pax2* and eya2 continued to be expressed in the OV after loss of Six1, consistent with a previous report in mouse that *pax2* otic expression does not depend upon Six1 (Ozaki et al., 2004). Second, the mouse OV data often predicts whether loss of Six1 will affect expression of target genes in other tissues. Although neural tube and retina expression of *pax2* also were normal in *six1*-null larvae, *eya2* expression in non-otic tissues was significantly reduced, suggesting that only in the OV can *eya2* expression be regulated independent of Six1. For the other 6 genes assayed, expression was reduced in OV as well as the other craniofacial tissues in which they are normally expressed.

### Craniofacial skeletal defects

In mouse, Six1 is required for craniofacial development, with loss of SIX1 causing eye, ear, lower and upper jaw defects (Laclef et al., 2003; Ozaki et al., 2004; Tavares et al., 2017). We found that in most Six1-heterozygote tadpoles (Figure 4A), all the cartilaginous elements of the tadpole head were indistinguishable from those of Six1 wild type tadpoles (data not shown), including the quadrate and Meckel’s cartilage, upper and lower jaw cartilages respectively and derived from the first branchial arch, the ceratohyal, a cartilage derived from the second branchial arch, and the branchial arch cartilage, derived from the posterior branchial arches (n=25). In contrast, in Six1-null tadpoles (Figure 4B), the quadrate, Meckel’s and the ceratohyal cartilages are hypoplastic and frequently deformed (Figure 4B; n=8). In addition, the otic capsule of Six1-null tadpoles (Figure 4B, D, n=8), a cartilage surrounding the entire inner ear, is significantly reduced in size when compared to the otic capsule of Six1-heterozygote tadpoles (Figure 4A, C, n=8). Table 2 shows the frequency of defects in Meckel’s, and ceratohyal cartilage, and otic capsule of Six1-null tadpoles. E14.5 Six1-null mouse embryos (Figure 4F) present similar defects in homologous craniofacial cartilages, including hypoplastic and deformed Meckel’s cartilage and severe hypoplasia of the otic capsule. Altogether, these results demonstrate that, as in mouse, loss of Six1 in *Xenopus* causes severe defects in the branchial arch-derived craniofacial cartilages.

**Table 2:** Frequency of dysmorphic cranial cartilages.

|  | **Meckel's** | **Ceratohyal** | **Otic capsule** |
| --- | --- | --- | --- |
| *six1*<sup>+/-</sup> | 8% (25) | 0% (25) | 0% (25) |
| *six1*<sup>-/-</sup> | 75% (8) | 87.5% (8) | 100% (8) |

**Figure 4:**
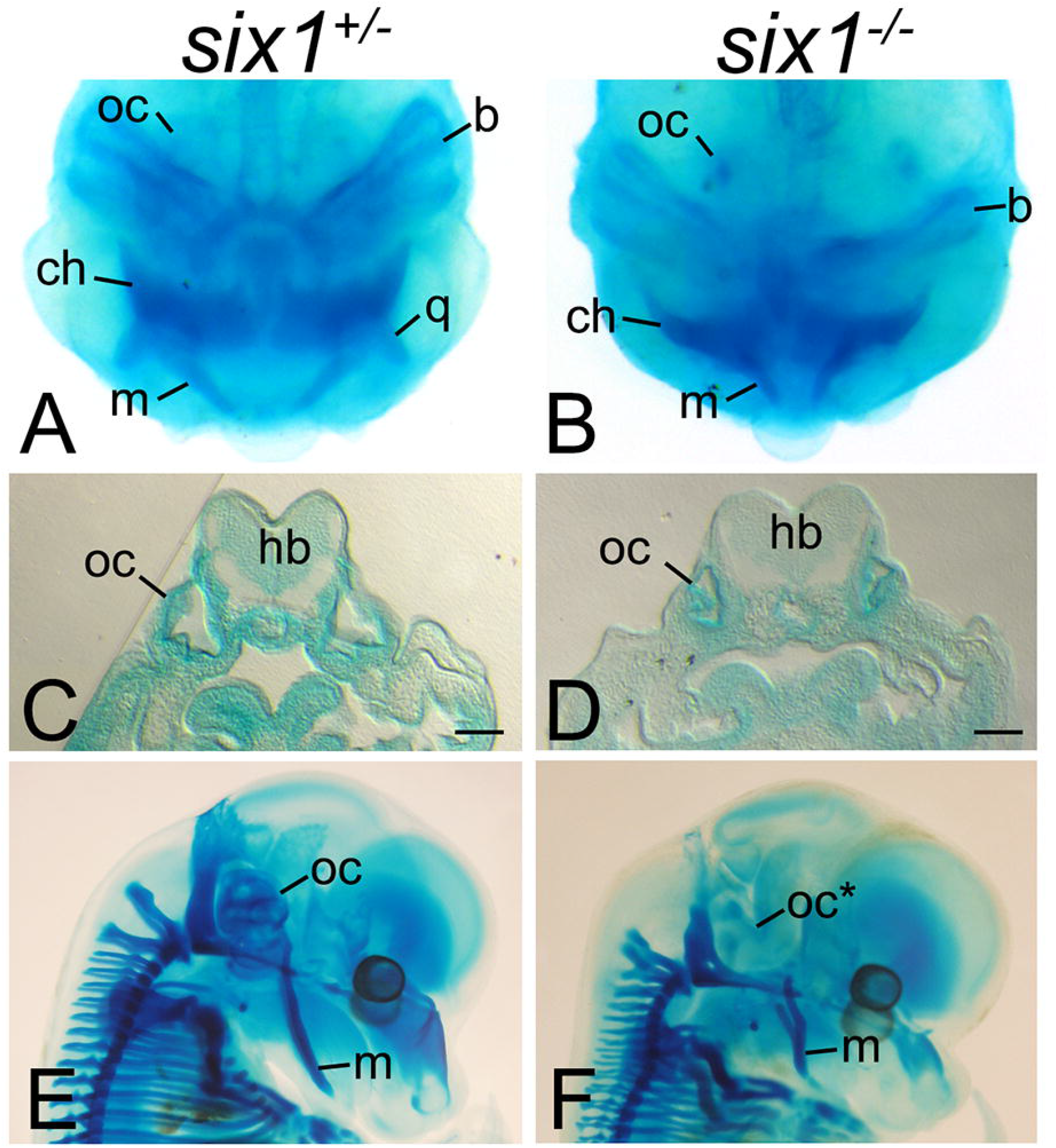
Loss of six1 causes severe craniofacial defects. A-B. Whole-mount Alcian blue staining of *Xenopus tropicalis* tadpoles shows that while *six1-*heterozygotes (*six1*^*+/-*^) present no detectable defects in the branchial arch-derived craniofacial cartilages, *six1*-nulls (*six1*^*-/-*^) have hypoplastic and deformed Meckel’s (m), quadrate (q), ceratohyal (ch), and branchial arch (b) cartilages and hypoplastic otic capsule (oc). C-D. Transverse section of tadpoles in A-B at the level of the otic capsule (oc) showing hypoplasia of this structure in Six1-null tadpoles (D). hb. hindbrain. Bar. 100μm. E-F. Whole-mount Alcian blue staining of E14.5 mouse embryos showing absent otic capsule (oc^*^) and deformed Meckel’s cartilage (m) in Six1-null embryos.

### Summary

This line of *Xenopus tropicalis* will be a useful resource for the community seeking to elucidate the role of Six1 in the development of several tissues, including olfactory sensory epithelium, inner ear, cranial sensory ganglia, kidney, branchial arches, and somitic and hypaxial muscle.

Since mutations in human *SIX1* underlie some cases of BOR and craniosynostosis (Smith, 2018; Calpena et al., 2021), this line also will be very useful for elucidating the role of this gene in these craniofacial syndromes.

## METHODS

### Mice

Generation and genotyping of *Six1*-null mice has been previously described (Ozaki et al., 2004). This line has been backcrossed onto the 129S6 background. The sex of embryos was not determined. All experiments were approved by the Institutional Animal Care and Use Committee (IACUC) at the George Washington University. Care of these mice followed local and national animal welfare law and guidelines. The George Washington University is certified by the Association for Assessment and Accreditation of Laboratory Animal Care.

### Crispr/Cas9 Editing

Two sgRNAs were designed to the first exon of *Xenopus tropicalis six1* using CRISPRScan (https://www.crisprscan.org/) (Moreno-Mateos et al, 2015), T1: <u>GG</u>TGAAGGCAAAGGCCGTGG and T2: <u>GG</u>GGCAGTGGGCAAGTACA; T1 target is located 150bp into exon 1, while T2 target is 312bp into exon 1. The GG nucleotides were added to the 5’end, for improved mutagenic activity (Gagnon et al 2014). sgRNAs were synthesized by in vitro transcription of the sgRNA PCR template using the T7 MEGAscript kit (Ambion, Cat. No. AM1334). F0 founders were produced by injecting T1 sgRNA into the eight vegetal blastomeres at the 32-cell stage; 0.25nl per blastomere with a total of 250pg of sgRNA and 500pg Cas9 per embryo. *X. tropicalis* Nigerian strain animals were used to generate this mutant line (RRID:NXR_1018). Seven F0 adults were outcrossed to wild type and F1 embryos were genotyped to assess germline transmission of mutations. Three of these founders (one male and two female) showed germline transmission. Both the +1/+ (*Xtr*.*six1*^*em1Horb*^, RRID:NXR_3089) and the -28/+ (*Xtr*.*six1*^*em2Horb*^, RRID:NXR_3145) *six1* mutants are available from the NXR (https://www.mbl.edu/xenopus).

### Husbandry and Genotyping of Xenopus

*Xenopus tropicalis* were housed in recirculating aquatic systems with established diet and water parameters (conductivity, pH, and temperature) as referenced in McNamara et al. (2018) and Shaidani et al. (2020). Embryos were generated by hormone induced natural mating of adult pairs of -28/+ *six1* mutants. Confirmed oocyte positive females were given 20 U of Pregnant Mare Serum Gonadotropin (PMSG) (Bio Vender, Cat. No. RP17827210000) and 80 µg of Ovine Luteinizing Hormone (oLH) (National Hormone and Peptide Program) (Wlizla et al., 2018), while virgin females were given 10 U of PMSG and 40 µg of oLH to prevent death via ovarian hyperstimulation syndrome (Green et al., 2007). Alternatively, 200 U of Human Chorionic Gonadotropin (hCG) (Bio Vender, Cat. No. RP17825010) can be used to induce ovulation in previously mated females and 100 U of hCG can be used for virgin females. Embryos were genotyped by collecting individual embryos and isolating genomic DNA using Sigma-Aldrich GenElute Mammalian Genomic DNA Miniprep Kit (G1N350-1KT). PCR amplification of the targeted region was done using the following primers: forward primer 5’-CCATGTCTATGCTGCCTTCC-3’ & reverse primer 5’-CCCTCAGTTTCTCTGCTTCC-3’.

PCR products were purified using NucleoSpin PCR Clean-up procedure (Macherey-Nagel 740609.250) and mutations were confirmed by sequencing. Adult frogs were genotyped by collecting genomic DNA from webbing of hindlimbs. Disposable biopsy punches were used to collect tissue (VWR 21909-140). F2 embryos were collected at desired stages and fixed in MEMFA (10 mL 10X MEMFA salts, 10 mL 37% formaldehyde, 80 mL NF H20); after fixation they were stored in Ethanol at -20C. For genotyping, genomic DNA was extracted from tailclips collected from fixed tadpoles after rehydration into PBS.

### In Situ Hybridization

*Xenopus* embryos were cultured to neural plate (st. 16-18) and larval (st. 28-34) stages (Nieuwkoop and Faber, 1994), fixed in 4% paraformaldehyde (PFA) and processed for *in situ* hybridization (ISH) as described previously (Yan et al., 2009). Antisense RNA probes were synthesized *in vitro* (MEGAscript kit; Ambion) as previously described (Yan et al., 2009). After analysis of whole embryos, a subset was embedded in a gelatin-based medium (0.5% gelatin, 30% bovine serum albumin, 20% sucrose, hardened with glutaraldehyde [75μl/ml]), and vibratome sectioned at 50 µm in the transverse plane. Whole-mount and serial section ISH images were collected with an Olympus SZX16 stereomicroscope coupled to an Olympus UC90 camera and cellSens Entry software.

### Cartilage Staining

Cartilage staining of *Xenopus tropicalis* tadpoles was performed according to Young et al. (2017). Tadpoles were fixed in 4% PFA for 1 hour at room temperature then incubated in a solution of acid/alcohol containing 0.1% Alcian Blue. When staining was complete, tadpoles were washed in the acid/alcohol solution without Alcian Blue, bleached with a solution containing 1.2% hydrogen peroxide and 5% formamide and cleared in 2% KOH with increasing concentrations of glycerol. Representative tadpoles were embedded in a gelatin-based medium (0.5% gelatin, 30% bovine serum albumin, 20% sucrose, hardened with glutaraldehyde (75μl/ml) and vibratome sectioned at 50μm in the transverse plane. Mouse embryos were harvested at E14.5, fixed in Bouin’s fixative, stained in 0.05% Alcian blue solution in 5% acetic acid, washed and clarified in benzyl alcohol:benzyl benzoate solution. Whole-mount and sectional images were captured using an Olympus SZH16 stereomicroscope coupled with an Olympus UC90 camera.

### Availability of mutant Xenopus lines

Both the +1/+ (*Xtr*.*six1*^*em1Horb*^, RRID:NXR_3089) and the -28/+ (*Xtr*.*six1*^*em2Horb*^, RRID:NXR_3145) *six1* mutants are available from the NXR (https://www.mbl.edu/xenopus).

## ACKNOWLEDGMENTS

Many of the methods were supported by Xenbase (http://www.xenbase.org/, RRID: SCR_003280) and the National *Xenopus* Resource (http://mbl.edu/xenopus/, RRID:SCR_013731). We thank Ms. Himani Datta Majumdar for performing ISH and sectioning embryos and tadpoles.

